# Neural Stimulation in vitro and in vivo by Photoacoustic Nanotransducers

**DOI:** 10.1101/2020.05.20.106898

**Authors:** Yimin Huang, Ying Jiang, Xuyi Luo, Jiayingzi Wu, Haonan Zong, Linli Shi, Ran Cheng, Shan Jiang, Xiaoting Jia, Jianguo Mei, Heng-Ye Man, Ji-Xin Cheng, Chen Yang

**Author notes:** Y. Huang and Y. Jiang contributed equally to this work. Corresponding author: Chen Yang.

## Abstract

Neuromodulation is an invaluable approach for study of neural circuits and clinical treatment of neurological diseases. Here, we report semiconducting polymer nanoparticles based photoacoustic nanotransducers (PANs) for neural stimulation. Our PANs strongly absorb light in the near-infrared second window and generate localized acoustic waves. PANs can also be surface-modified to selectively bind onto neurons. PAN-mediated activation of primary neurons *in vitro* is achieved with ten 3-nanosecond laser pulses at 1030 nm over a 3 millisecond duration. *In vivo* neural modulation of mouse brain activities and motor activities is demonstrated by PANs directly injected into brain cortex. With millisecond-scale temporal resolution, sub-millimeter spatial resolution and negligible heat deposition, PAN stimulation is a new non-genetic method for precise control of neuronal activities, opening potentials in non-invasive brain modulation.

## Introduction

Neural stimulation is an important tool enabling our understanding of how brains function and treating neurological disorders. Electrical stimulation is the basis of current implantable devices and has already used in the clinical treatment of depression, Parkinson’s and Alzheimer’s diseases. These devices, often made of metal electrodes, are limited by their invasive nature ^1^, inability to targeting precisely due to current spread, and its MRI incompatibility. Noninvasive clinical or pre-clinical methods, such as transcranial magnetic stimulation (TMS) ^2^, transcranial direct current stimulation (tDCS) ^3^ and focused ultrasound ^4,5^, do not require a surgical procedure but offer a spatial resolution on the order of several millimeters. Optogenetics has been shown as a powerful method modulating population neural activities in rodents more precisely and with cell specificity ^6,7^. Optogenetics requires genetic modification through viral infection, which makes it challenging to be applied to humans ^8^.

Nanostructures target neuron membrane locally, convert and amplify the external excitation to local stimuli, offering new interfaces as promising alternative neural stimulation approaches. Gold nanoparticles and nanorods were studied for photothermal neural stimulation in vitro ^9-12^. Gold nanoparticles and carbon nanotubes were also used for photothermal-driven optocapacitive stimulation in vitro ^13-15^. The Tian and Bezanilla groups reported photoelectrical stimulations with silicon nanostructures ^16^. The light wavelengths used in these light driven stimulations were mostly in the range of 520 – 808 nm, which has limited penetration through skulls and in brain tissue. To offer deeper penetration, thermal stimulation triggered by nanoparticles absorbing longer wavelength light or magnetic field has also been investigated. The Pu group demonstrated photothermal neural stimulation in vitro using bioconjugated polymer nanoparticles absorbing 808 nm and binding to transient receptor potential cation channel subfamily V member 1 (TRPV1) ^17^. The Anikeeva group used gene transfection to over-express the thermal sensitive ion channels and then utilized the magneto-thermal effect of the paramagnetic nanoparticles to activate these channels ^18^. In these studies, significant local temperature rise, exceeding the thermal threshold of the ion channels, e.g. 43 °C in the case of TRPV 1, was observed, thus raising concerns over safety of thermally activated brain stimulation. The Khizroev group used the magneto-electric nanoparticles under an applied magnetic field to perturb the voltage-sensitive ion channels for neuron modulation ^19^. Notably, these magnetic stimuli-based techniques deliver a spatial precision relying on the confinement of the magnetic field, which is on the millimeter to centimeter scale. New technologies and concepts are still sought to achieve non-invasive, genetic free and precise neural stimulation.

Here, we report the development and application of photoacoustic nanotransducers (PANs) to enable non-genetic neural stimulation in cultured primary neurons and in live brain (**Figure 1a**). Our PANs, based on synthesized semiconducting polymer nanoparticles, efficiently generate localized ultrasound by an optoacoustic process upon absorption of nanosecond pulsed light in the NIR-II window (1000 nm to 1700 nm) (**Figure 1b)**. The NIR-II light has the capability of centimeter-deep tissue penetration ^20,21^, which is beyond the reach of visible light currently used in optogenetics. We further modified the PAN surface for non-specific binding to neuronal membrane and specific targeting of mechanosensitive ion channels, respectively. We showed that upon excitation at 1030 nm PANs on the neuronal membrane successfully activated rat cortical neurons, confirmed by real time fluorescence imaging of GCaMP. We then demonstrated *in vivo* motor cortex activation and invoked subsequent motor responses through PANs directly injected into a mouse living brain. Importantly, the heat generated by the ns laser pulses is confined inside the PAN, resulting in a transient temperature rise during the photoacoustic process, evident by COMSOL simulations. Collectively, our finding shows photoacoustic nanotransducers as a new platform for modulating neuronal activities. It is triggered by NIR II light and shows neglectable temperature increase, opening up opportunities for deep-penetrated-light controlled neural activation with high precision.

**Figure 1.**
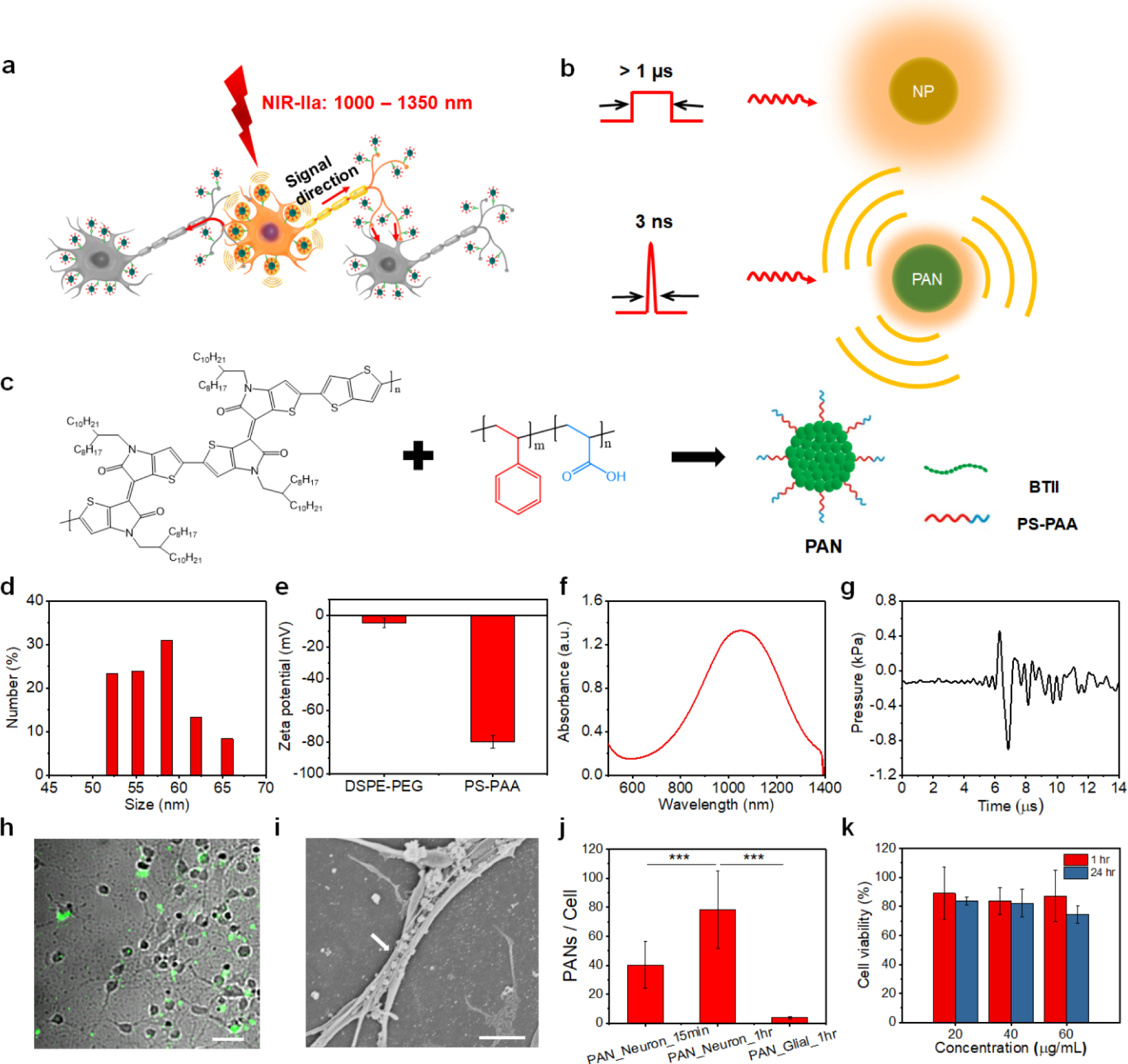
Surface modified PANs sufficiently bind to neurons. (a) Scheme of the PAN induced neural stimulation. (b) Schematic illustrating that upon nanosecond laser pulse excitation a PAN generates photoacoustic signals. NP: nanoparticle. (c) Schematic of PAN synthesis. (d) Dynamic light scattering analysis of PAN solutions. (e) Zeta potential measurement of PAN solutions with a concentration of 1.0 mg/mL with DSPE-PEG and PS-PAA functionalization, respectively. (f) UV-Vis spectrum of PAN solution with a concentration of 1.0 mg/mL. (g) Photoacoustic signal measured from PAN solution (1.0 mg/mL). (h) TA images of PANs binding to neurons after 15-minute culture. Scale bar: 50 μm. (i) SEM image of PAN binding to neurons. Scale bar: 2 μm. (j) Binding density analysis of PANs to soma region of neurons and glial cells at 15 minutes and 1 hour, respectively. (n.s.: non-significant, p > 0.5; *: p < 0.5; **: p < 0.01; ***: p < 0.001) (k) Cytotoxicity analysis of PANs to neurons by MTT assay.

## Results

### Synthesis of PANs

We first synthesized NIR-II absorbing semiconducting polymer bis-isoindigo-based polymer (BTII) ^22^. To obtain nanoparticles, we then modified the polymer with polystyrene-block-poly(acryl acid) (PS-b-PAA) via a nanoprecipitation method (**Figure 1c**). The PS-b-PAA was chosen due to the amphiphilic nature of its chemical structure. The hydrophobic polystyrene portion forms a π-π stacking with the polymer, while the hydrophilic poly(acryl acid) (PAA) makes the polymer into water-soluble nanoparticles with carboxyl groups decorated on the surface. FT-IR spectrum confirmed the presence of carboxyl groups (**Figure S1**), indicating the successful modification. The size of nanoparticles prepared was measured to be 58.0 ± 5.2 nm using dynamic light scattering (DLS) (**Figure 1d**). The nanoparticles were found to be negatively charged indicated by a potential of -79.79 ± 4.04 mV through the zeta potential measurement. This negative charge is attributed to PS-b-PAA surface modification, as we found that a neutrally charged surfactant 1,2-distearoyl-sn-glycero-3-phosphoethanolamine-N-[(polyethylene glycol)-2000] (DSPE-PEG(2000)) modified PANs are charged with - 4.88 ± 3.06 mV (**Figure 1e**).

### PANs generate strong acoustic waves under NIR-II light pulses

The planar backbone of the semiconducting polymer chain pushed the absorption to the NIR-II window ^23^. We confirmed this by Ultraviolet (UV)-Visible-NIR spectroscopy. The nanoparticles absorb broadly NIR-II light from 800 to 1300 nm with a peak at 1100 nm (**Figure 1f**). Measured with an ultrasound transducer with a central frequency at 5 MHz, 1.0 mg/mL nanoparticle solution exhibits a photoacoustic signal showing a waveform in time domain with approximately 2 μs in width and a peak to peak amplitude of 33.95 mV (**Figure 1g**), under 1030 nm nanosecond laser with a pulse width of 3 ns, a repetition rate of 3.3 kHz, and an energy density of 21 mJ/cm^2^. The peak pressure was estimated to be 1.36 kPa. Since these nanoparticles generate a strong photoacoustic signal under pulsed NIR-II light, we termed them photoacoustic nanotransducers (PANs) and studied their potential for neural binding and stimulation, as detailed below.

### PANs sufficiently bind to neurons

As recently reported, nanoparticles with negatively charged surface can bind onto neuronal membrane, whereas positive and neutral nanostructures showed no interactions with neurons ^24^. To examine whether negatively charged PANs can bind onto the neuron membrane, we cultured PANs with embryonic cortical neurons collected from Sprague Dawley (SD) rats. The neurons were first cultured for 15-18 days (Days in vitro, DIV15-18). We then added 150 μL 20 μg/mL PAN solution into the culture, reaching a concentration of 2 μg/mL. The same concentration was used in all experiments in this work otherwise noted.

Being able to quantify and control the binding density of PANs to neurons is critical for successful stimulation. Since the semiconducting polymer show strong intensive intrinsic transient absorption (TA) signals, we then used label free TA microscopy to visualize binding of PANs on neurons ^25,26^. Specifically, we used 200 fs laser pulses at 845 nm and 1045 nm as the pump and probe beams, respectively, with laser power fixed at 20 mW for both beams for TA imaging. To quantify the effective density of PANs bound to neurons, we first measured the signal-to-noise ratio (SNR) of the TA signals of PAN solutions with concentrations ranging from 2.0 to 55.0 μg/mL to obtain a TA calibration curve. The SNR of TA signals was found to be linear to the PAN concentration with a slope of 14.24 mL/μg. Next, we incubated neurons in culture supplemented with PANs for 15 minutes, rinsed three times with PBS to remove unbound PANs, and fixed the cells for TA imaging. The PANs were found to bind onto the neurons at an estimated density of 40.2 ± 15.9 PANs per soma (**Figure 1h**). The number of PAN was calculated based on effective TA concentration estimated based on the measured TA intensity and the TA calibration curve, focused spot volume, and estimated molecular weight of PANs. Scanning electron microscopy (SEM) images also confirmed the presence of PANs in the embryonic neuron cells in both neurite (**Figure 1i**) and soma regions (**Figure S3**), respectively. By increasing the culture time to 1 hour, a higher binding density was achieved and the number of PANs per neuron on the soma area was found to be 78.1 ± 26.7 (**Figure 1j** and **Figure S4a**). As a comparison, no PANs bind to the glial cells after 1-hour culture (**Figure S4b**).

To test the cytotoxicity, MTT assay was performed on cultured neurons following incubation with PANs for 1 hour and 24 hours, respectively. Cell viabilities over 80% were observed in all experimental groups with PAN concentrations ranging from 20 to 60 μg/mL (**Figure 1k**), indicating low toxicity of PANs to neurons. These results collectively show that negatively charged PANs can sufficiently bind onto neuronal membranes via a charge-charge interaction, without showing obvious cytotoxicity.

### PANs stimulate primary neurons in culture

After showing that PANs bind to neurons, we further investigated their potential for neural stimulation. We performed calcium imaging on Sprague Dawley (SD) rat primary cortical neurons transfected with GCaMP6f. Neuron cultures were transfected with GCaMP6f at DIV 5, and at DIV 15-18 were incubated with 150 μL, 20 μg/mL PAN solution for 15 minutes. Fluorescence imaging was performed with a wide field fluorescence microscope at the excitation wavelength of 470 nm. Nanosecond laser at 1030 nm was delivered to the culture via a 200 μm diameter optical fiber with 0.22 NA. Conditions of the pulsed laser include a pulse width of 3 ns, a repetition rate of 3.3 kHz, a laser train of 3 ms (corresponding to 10 laser pulses), and an energy density of 21 mJ/cm^2^. Fluorescence imaging was performed on 5 culture batches for each group. Data from total 60 neurons, all of which were within 100 μm proximity to the surface of the fiber were analyzed. 100 μm was chosen based on the estimated illumination area of the optical fiber. A representative fluorescence image of the neuron culture is shown in **Figure 2a**, with the dashed circle showing the position of the fiber. Increase in fluorescence intensity of GCaMP6f at individual neurons was clearly observed immediately after applying pulsed laser, showing in the real-time video (**Movie S1**). Out of total 60 neurons studied, 51 neurons showed an increase in fluorescence greater than 10% or *F/F*_*0*_ ratio above 1.10 after the laser onset (**Figure 2b**). *F*_*0*_ is the baseline fluorescence signal of the neurons before the stimulation.

**Figure 2.**
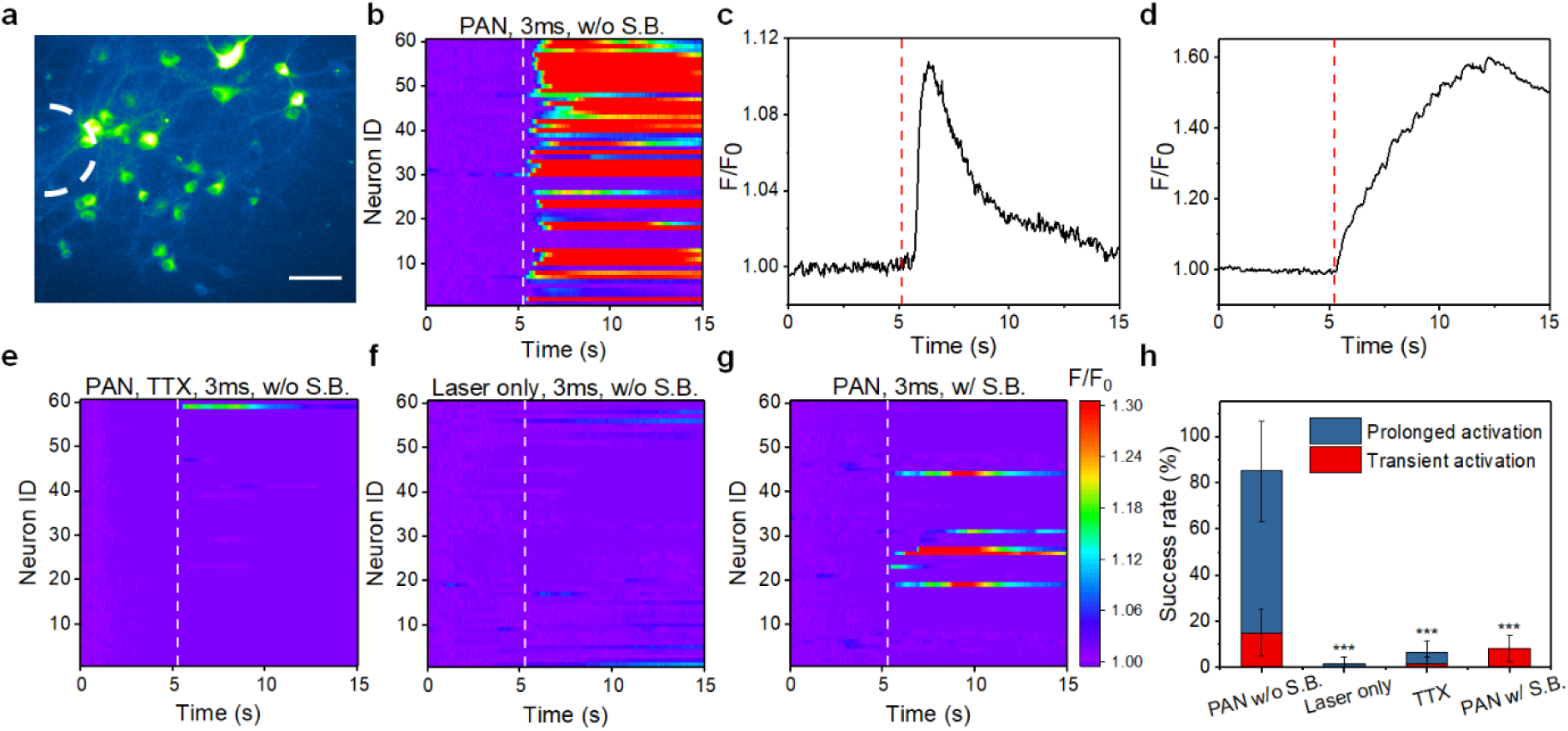
PANs induce neural stimulation with over an 80% success rate. (a) Representative fluorescence image of GCaMP6f labeled neurons (DIV 15-18) cultured with PANs for 15 minutes. Scale bar: 100 μm. White dash line indicates the position of the optical fiber delivering ns pulsed light. (b) Colormaps of fluorescence changes of neurons stimulated by PANs using the 1030 nm nanosecond laser with a 3 ms laser pulse train. White dashed lines indicate laser is on. (c,d) Representative fluorescence changes as a function of time for transient activation (c) and prolonged activation (d), respectively (N=60). Red dashed lines indicate laser is on. (e-g) Colormaps of fluorescence changes of neurons treated with laser only (e), with TTX added into the culture medium (f) and with the synaptic blocker cocktail added in the culture medium. Same laser conditions were used as (c) and (d). All Colormaps were plotted under the same dynamic range. (h) Success rate analysis. Error bars: ±SD. (p value was calculated using PAN w/o S.B. group as reference. n.s.: non-significant, p > 0.5; *: p < 0.5; **: p < 0.01; ***: p < 0.001)

Notably, two types of responses were detected, a transient response shown in **Figure 2c** and a prolonged response taking longer time (up to 60 s) to recover to the baseline shown in **Figure 2d** and **Figure S5**. We fitted the decay of the response curves exponentially and defined a time constant when they decrease by a factor of *1/e* (0.368) from the peak fluorescence intensity. The transient activations typically show decay time constants ranging from 2 to 5 s, while the prolonged activations have time constants ranging in 5 to 10 s (**Figure S6**).

A success rate, defined as the percentage of activated neurons identified through the *F/F*_*0*_ ratio above 1.10, was calculated. Under the 3 ms laser pulse train, total 85.0% of the neurons exhibited activations immediately after the nanosecond laser was onset. Specifically, 15.0 ± 10.0% and 70.0 ± 21.8 % were observed as the transient responses and prolonged responses, respectively (**Figure 2h**).

To investigate whether the activations observed based on the increased fluorescence intensity are caused by action potential activation, we performed a control with addition of 3 μM of Tetrodotoxin (TTX), a block of voltage-gated sodium channels. After addition of TTX, only a total of 6.7% neurons showed activation, with 1.7 ± 2.9% for transient activation and 5.0 ± 5.0 % for prolonged activation (**Figure 2e**). As an additional control, only applying nanosecond laser at the same laser condition without PANs was performed. Only 1.7 ± 2.9% neurons showed prolonged response and no transient response was observed, indicating optical excitation through ns laser alone triggers negligible activities (**Figure 2f**). We further applied a cocktail of synaptic receptor blockers, 10 μM NBQX, 10 μM Gabazine and 50 μM DL-AP5 ^27^, to study the networking effect on activation. As shown in **Figure 2g**, application of synaptic blockers significantly reduced the success rate of transient activation to 8.3 ± 5.8% and blocked the prolonged activation completely. These results show that observed activations without synaptic blockers are largely contributed by synaptic inputs from a subpopulation of neurons directly stimulated initially by PANs, whereas the observed prolonged response is likely due to sustained activities within the neuronal network. Notably, no direct activation was found outside the illumination area of the optical fiber (**Figure S7**), indicating the stimulation area is defined by the illumination or focus of the pulsed light and a spatial confinement of a few hundred microns is achieved in vitro using the optical fiber. Collectively, we have confirmed that the observed PAN-triggered neuroactivities are action potential-dependent and require activation of neurotransmitter receptors.

Key parameters to control the stimulation through PANs include laser conditions and binding density of PANs on neurons. To understand the effect of the pulsed laser train on activations by PANs, we first studied the activation under increased laser train of 5 and 10 ms, corresponding to 17 and 33 laser pulses, respectively. In the laser only groups, the overall success rate was found to be 3.3 ± 2.0% using 5 ms, and 18.3 ± 10.4% for 10 ms (N=60, 3 different culture batches), dominated by the prolonged activation (**Figure S8 a-c**). With the presence of PANs after a 15 min culture with neurons, under the 5 ms laser duration, an overall success rate of 66.7 ± 14.4% was observed (N=60, 3 different culture batches). When the laser pulse train increased to 10 ms, the total success rate was found to be 80.0 ± 15.3 % (**Figure S8 d-f**). Notably, both 5 ms and 10 ms laser pulse trains produced neural activities dominated by prolonged activation. Therefore, 3 ms was found as a sufficient pulse train to produce a high successful rate in direct activation with a less networking effect. Therefore, we identified it as the optimal laser pulse train for PAN mediated neural stimulation given other laser parameters used.

To investigate how the binding density impacts PAN mediated stimulation, we varied the incubation time of PANs with neuron cultures. In the group where the stimulation was performed immediately after addition of PANs followed with rinses, no neural activation was detected (**Figure S9a**). This observation confirmed that only bound PANs can trigger the activation. In the group where the stimulation was performed after PANs were incubated with neurons for 1 hour, 20.0 ± 18.0 % neurons exhibited transient activations and 28.33 ± 16.07 % exhibited prolonged activation (**Figure S9b** and **c**). These results indicated 15-minute culture time provides a binding density sufficient to trigger neural stimulation.

### PANs conjugated with TRPV4 enable channel-specific neural stimulation

To enable specific targeting for stimulation, we bioconjugated the PANs with antibodies to specifically target the mechanosensitive ion channel transient receptor potential cation channel subfamily V member 4 (TRPV4). TRPV4 was chosen based on its high expression rate on the neuronal cell membranes and its capability in sensing external mechanical stimuli ^28,29^. We conjugated PANs with anti-TRPV4 antibody through a carbodiimide coupling reaction, using ethyl(dimethylaminopropyl) carbodiimide (EDC) and N-hydroxysuccinimide (NHS), between the carboxyl group on PANs to the amine group on the antibody (**Figure 3a**) ^17^. Successful bioconjugation was confirmed by comparing characteristics of PANs and anti-TRPV4 conjugated PANs (PANs-TRPV4). Electrophoresis showed that PANs-TRPV4 migrated much less compared with PANs (**Figure 3b**). Additionally, a size increase from 59.4 ± 5.3 nm to 181.8 ± 36.7 nm was revealed by DLS analysis (**Figure 3c**). The zeta potential for PAN-TRPV4 is -1.49 ± 0.38 mV, almost neural (**Figure 3d**). No color change was noticed in the PAN-TRPV4 solution (**Figure S10a**). No obvious shift in absorption spectrum was identified for the PAN-TRPV4 solution (**Figure S10b**).

**Figure 3.**
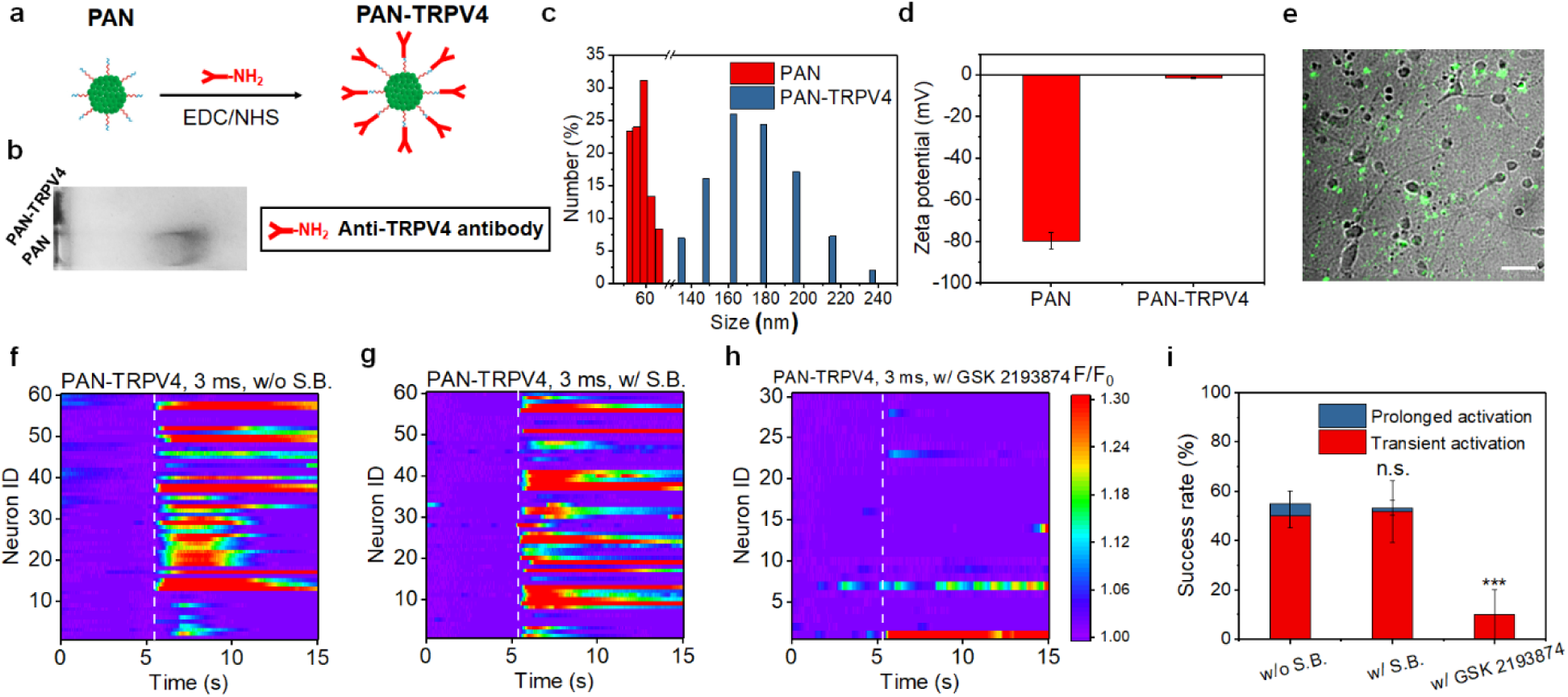
PANs-TRPV4 induce transient activation of neurons. (a) Schematic of PAN-TRPV4 synthesis. (b) Electrophoresis measurements of PAN and PAN-TRPV4 solutions with a concentration of 0.1 mg/mL. (c) DLS analysis of PAN and PAN-TRPV4 sizes. (d) Zeta potential measurement of PAN and PAN-TRPV4 solutions. (e) TA image of PANs-TRPV4 cultured with neurons for 15 minutes. Scale bar: 50 μm. (f-h) Colormaps of fluorescence intensity change in neurons treated with PANs-TRPV4 without synaptic blockers (f), with the synaptic blocker cocktail (g), with GSK 2193874 (h) added in the culture medium. A laser pulse train of 3 ms was used in all experiments. (i) Success rate analysis. Error bars: ±SD. (p value was calculated using w/o S.B. group as reference. n.s.: non-significant, p > 0.5; *: p < 0.5; **: p < 0.01; ***: p < 0.001)

We then confirmed the expression of the TRPV4 channels in the membrane of embryonic cortical neurons. As indicated in IF staining images (**Figure S11)**, TRPV4 channels are expressed vigorously throughout the soma and neurites of the neurons, as expected ^30,31^. This result indicates that a high number of target sites on the neuronal membrane are available for PANs-TRPV4 for potential binding. After incubation with PANs-TRPV4 for 15 minutes under the same condition as for PANs, PANs-TRPV4 binding to neurons were visualized by TA microscopy (**Figure 3e)**. The PAN-TRPV4 density was estimated to be 43.8 ± 20.8 per soma, slightly larger than that found for PAN binding.

Next, we studied whether the PAN-TRPV4 could improve the performance of neural stimulation through direct binding to the mechanosensitive ion channels. Under the same experimental condition used for PANs, we analyzed 60 neurons collected from 5 different culture batches to statistically quantify the neural stimulation. As shown in **Figure 3f** and **Movie S2**, neural activations induced by PAN-TRPV4 show an overall success rate of 55.0%, of which the transient stimulation responses is 50.0 ± 5.0% and the prolonged response is 5.0 ± 5.0 % (**Figure S12**). The portion of transient activation increased substantially, compared to that for PANs. As shown in **Figure 3g**, with synaptic blockers in the culture medium, the overall success rate remains as 53.3%. 51.7 ± 12.6 % of neurons showed transient activation and only 1.7 ± 2.9 % showed prolonged activation, which indicates the transient activation was not blocked by synaptic blockers and that direct activation was achieved using PANs-TRPV4. To further validate that the observed activation is mediated by the activation of the TRPV4 channel, we performed a control with adding a specific TRPV4 channel blocker, GSK 2193874 ^32^, into the culture, prior to adding PAN-TRPV4 solution (N=30, collected from 3 different culture batches). As shown in **Figure 3h**, with the presence of TRPV4 channel blockers, the success rate significantly decreased, with 10.0 ± 10.0% of the neurons showing transient response and no prolonged activation was detected (**Figure 3i**). These results collectively confirm that PAN-TRPV4s enabled a specific stimulation directly through the TRPV4 ion channel.

### *In vivo* neural stimulation by PANs

Upon successful stimulation of cultured primary neurons, we asked whether PANs could activate neurons in vivo in living animals. We directly injected 600 nL of 1.0 mg/mL PAN solution into the primary motor cortex of C57BL/6 mice using a stereotaxic injector at a controlled speed of 20 nL/min. Stimulation was performed 1 hour after the injection. To validate brain activation, we performed local field potential (LFP) recording at the PAN injection site. To avoid electric artifact produced by laser radiation on the metal electrode, we used multifunctional fibers with a thick polymer coating as the recording electrodes (**Figure S13)** ^33,34^. As shown in the **Figure 4a** and **b**, 3 ms laser pulse train at 27 mJ/cm^2^ produced strong LFP response on the stimulated cortex, while in the control group on the contralateral side without PAN injection, the laser irradiation did not produce any response.

**Figure 4.**
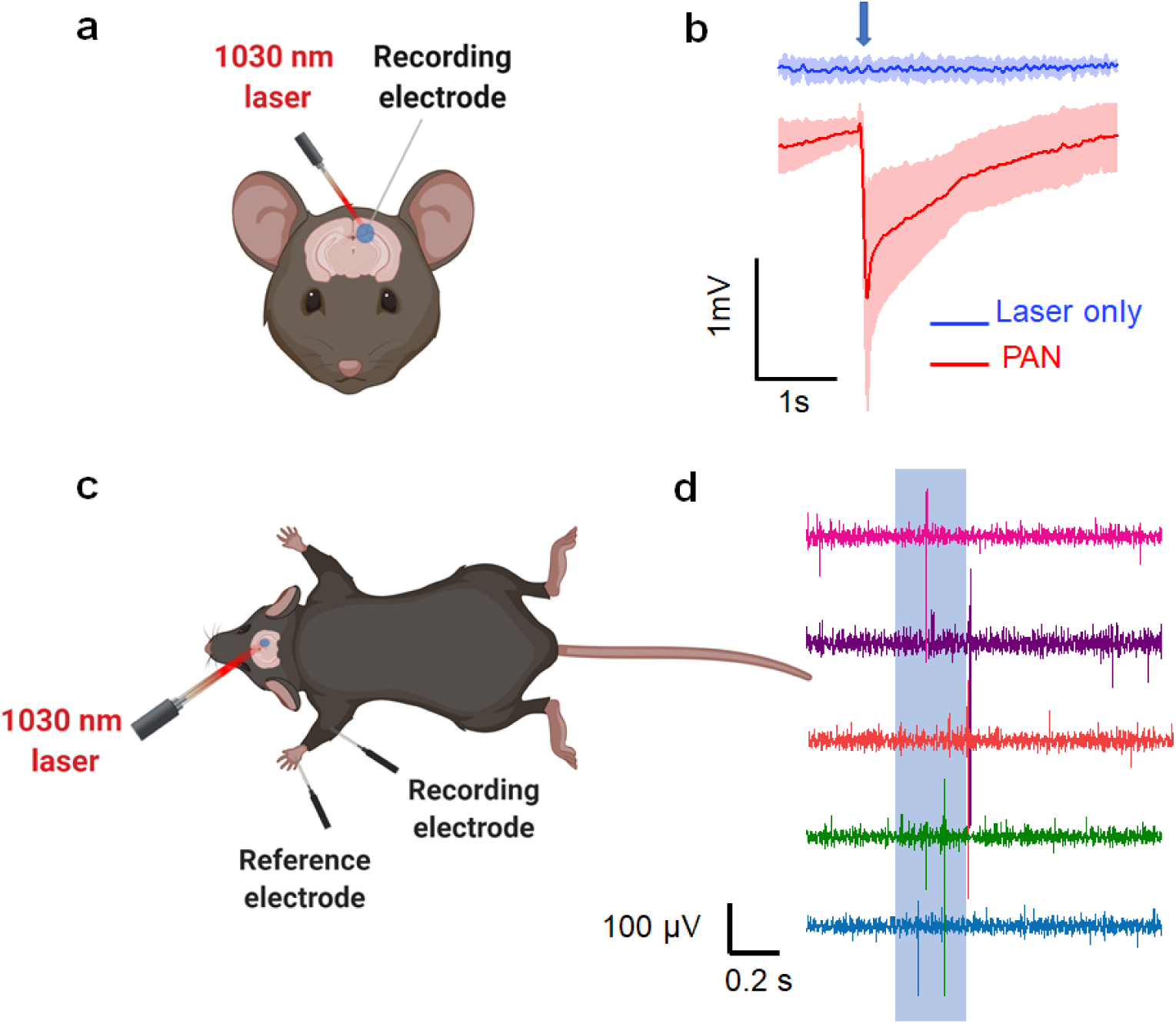
In vivo neural stimulation by injected PANs. (a) Schematic of in vivo electrophysiology measurement. (b) Representative electrophysiology curves measured at the brain region without PANs as the control group (blue) and PAN treated region (red) on 3 mice. Blue arrow indicates laser is on. (c) Schematic of EMG recording. (d) Representative curves of EMG measurements. All schematic diagrams were created using Biorender.

With successful LFP recording of PAN stimulation in brain, we further evaluated the behavior outcome of the stimulation. We performed electromyography (EMG) as a measurement of the effect of PAN brain stimulation. 600 nL PAN solution at 1.0 mg/mL was injected to the primary motor cortex of the mouse. At 1 hour after the injection, a needle electrode was inserted subcutaneously and parallel to the forelimb triceps brachii muscle. A reference electrode was inserted in the footpad with a ground electrode inserted subcutaneously on the trunk and ipsilateral to the stimulation site (**Figure 4c**). A 200 ms laser pulse train was delivered to the injection site through an optical fiber. EMG responses with an amplitude of 428.8 ± 119.0 μV, with a delay of 127.8 ± 24.3 ms, were recorded (**Figure 4d**). These results suggest that the PAN mediated brain stimulation was sufficient to induce motor cortex activation and invoke subsequent motor responses.

### PAN-mediated stimulation is not thermally induced

The photoacoustic effect is known to associated with a temperature increase. To gain insights on how much the photothermal process might contribute to the successful activation discussed above, we performed neuron stimulation under continuous wave laser (CW). CW laser excitation of nanoparticles is known to produce a local temperature rise without generation of photoacoustic signals ^35^. By comparing neural response to PANs upon excitation by the CW laser to that upon excitation by the nanosecond laser at the same power, we can determine how different our stimulation is from the known nanoparticle mediated photothermal stimulation. Since PANs absorb broadly in the range of 800 to 1300 nm, we used a CW laser at 1064 nm. Identical neuronal culture conditions were used.

While we achieved successful neural activation under ns laser power of 61 W/cm^2^ and a train of 10 pulses over 3 ms, no activation was detected using CW laser excitation with the laser power of 61 W/cm^2^ over 3.9 ms duration (N=30, 3 different cell culture batches, **Figure 5a**). No activation was observed as we increased the CW laser power to 397 W/cm^2^ while maintaining the CW laser duration at 3.9 ms (N=30, 3 different cell culture batches, **Figure 5b**). Activation of neurons was only observed when the duration was increased to 2.5 s at 397 W/cm^2^ (N=20, 3 different cell culture batches, **Figure S14**). These results show that under the CW laser at compatible power and duration to nanosecond laser conditions, the photothermal effect produced by the PANs alone cannot result in neural activation. The photoacoustic function of PANs enabled by the nanosecond light pulse contributed dominantly to the activation.

**Figure 5.**
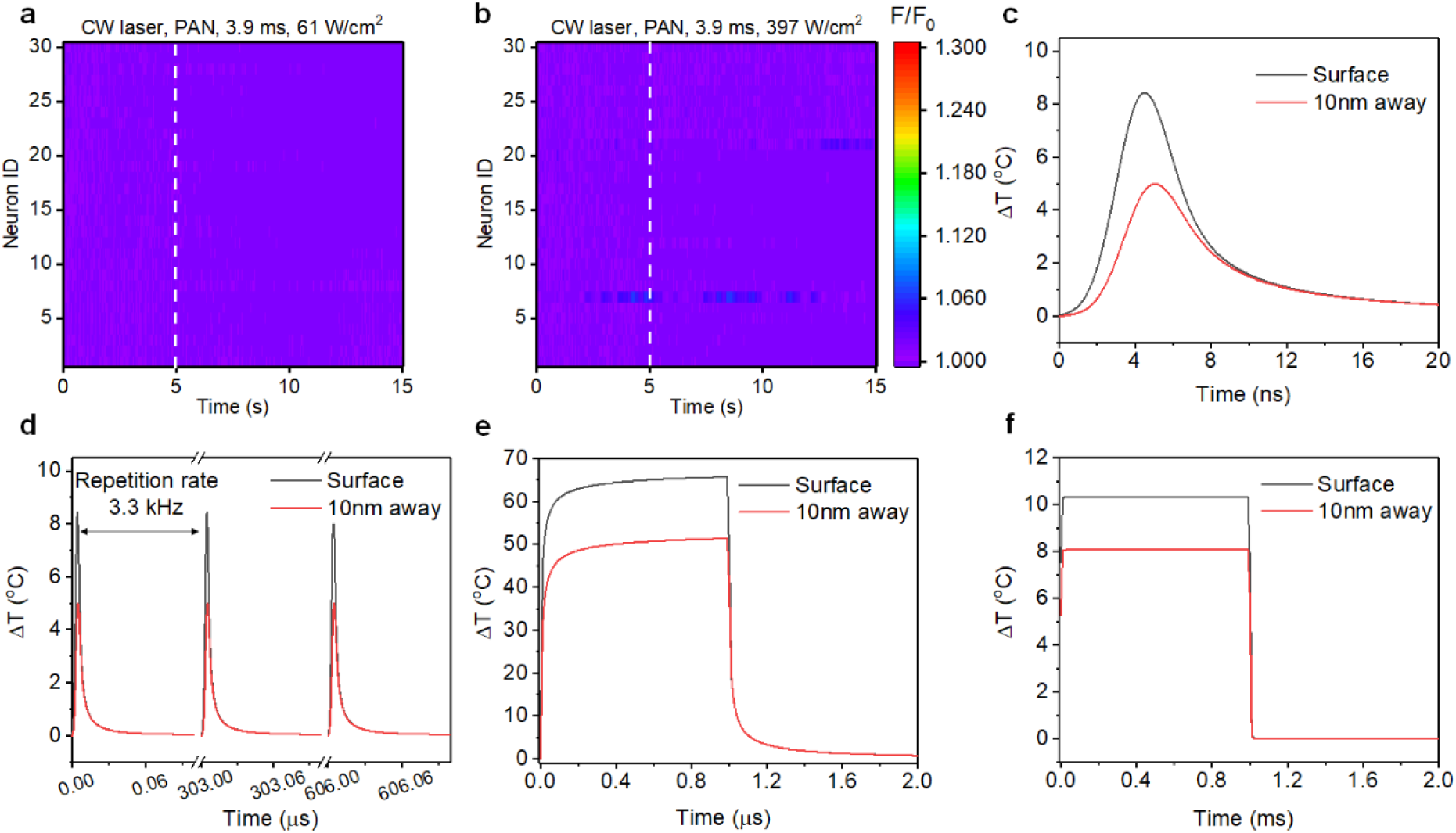
PAN-mediated neural stimulation is not thermally induced. (a,b) Colormaps of fluorescence intensity change of neurons cultured with PANs under a 1064 nm CW laser with laser condition of (a) a 3.9 ms laser duration and power density of 61 W/cm^2^; (b) 3.9 ms laser duration and power density of 397 W/cm^2^. All Colormaps were plotted under same dynamic range. Neurons were cultured with PANs for 15 min before stimulation. (c, d) COMSOL simulation on temperature changes with a single 3 ns laser pulse (c) and three 3 ns laser pulses with a 3.3 kHz repetition rate (d) on PAN surface (black) and 10 nm away from PAN surface in water (red). Laser wavelength is 1030 nm. Pulse energy is 1.18 nJ. (e, f) COMSOL simulation on temperature changes for a gold nanoparticle with 1 µs laser and pulse energy of 67.8 nJ, and (f) gold nanoparticle with 1 ms laser with pulse energy of 9.8 µJ. Laser wavelength is 532 nm.

To understand how temperature rises and dissipates upon ns laser excitation of a nanoparticle, we applied COMSOL to simulate the evolution of PAN surface temperature in water. Simulation for temperature at 10 nm away from surface of PAN in water was also performed, aiming to probe the temperature of neuron membrane where a PAN binds to. **Figure 5c** shows how the PAN temperature evolves under excitation by a single 3-ns laser pulse at 1030 nm. Pulse energy of 1.18 nJ was used accordingly to the condition used in our PAN stimulation experiments. Temperature increase is found to quickly rise to a peak value of 8.4 °C on the PAN surface (black, **Figure 5c**) and to 5.0 °C at 10 nm away from the PAN surface (red, **Figure 5c**), respectively. Importantly, in both cases, temperature decays to the baseline within 10 nanoseconds from the peak value. We note that the laser pulse train used for PAN stimulation is operated with a repetition rate of 3.3 kHz. At this repetition rate, the laser pulse train resulted in pulsed temperature spikes with a FWHM of 3 nanoseconds and no temperature accumulation is found (**Fig. 5d**). To compare, we simulated the temperature evolution for gold nanoparticles of 60 nm diameter under 532 nm CW laser with conditions reported for successful photothermal driven optocapacitive stimulation. Two conditions, one with energy of 38.9 nJ and duration of 1 µs and the other with energy of 7.41 µJ and duration of 1 ms, respectively, were used (**Fig. 5e and f**) ^13^. The temperature evolution under CW laser excitation was found to be substantially different from the temperature evolution under the ns laser excitation. As shown in **Figure 5e** and **5f**, the temperature increases on the Au nanoparticle surface quickly reach a plateau within the first 200 ns in both CW cases, with plateaued values at 65.6 °C and 10.3 °C, respectively. Similar temperature features also found in our simulation of graphite microparticles under a laser energy of 0.7 µJ for 80 µs laser duration at 532 nm, consistent with reported experimental and calculation results ^14,15^. These CW laser conditions were reported to generate a photothermal induced optocapacitive neuron stimulation, in which 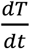 is identified as a drive for 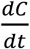, resulting in a current leading to action potential activation. Our results clearly demonstrate the difference in 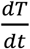 between the PAN under the ns laser and reported nanoparticles under CW laser. In addition, in the PAN case, the duration of each temperature spikes is a few nanoseconds, more than 100 times smaller than that found for nanoparticle under CW. It is conceivable that current induced by capacitance change over these tens of nanoseconds can be negligible. Together, our results suggest that the PAN stimulation is distinct from the photothermal optocapacitive stimulation.

## Discussion

In this work, we demonstrated semiconducting polymer-based PANs for neural stimulation under excitation by a nanosecond laser at NIR-II window. A over 80% success rate was achieved in vitro. Enhanced specificity was demonstrated via bioconjugating TRPV4 to the PANs. Successful in vivo activation through PANs directly injected into the cortex area of mouse living brains was demonstrated by LFP and EMG recording.

The photothermal effect of nanoparticles has been reported to successfully modulate neurons mainly in vitro ^9,11-13,17,36-40^. Two potential stimulation mechanisms were proposed, one through the increase of temperature, with highest temperature often found in the range of 50 °C to 70 °C ^17,38^, and another through an optocapacitive stimulation determined by the rate of temperature change ^13-15^. In our work, excited by a 3-nanosecond pulsed laser, the maximum temperature rise on the PAN surface is 8 °C and temperature change is in the form of 10 spikes, each of which is less than 10 nanoseconds in duration, over 3 ms. It is conceivable that such transient temperature spike would not result in current flow through a neuronal membrane. Instead, the PANs are able to generate a localized acoustic wave on the microsecond scale upon ns pulsed light (**Figure 1**). Activation did not occur when we changed the ns laser to a CW laser of the same energy (**Figure 4a** and **4b**). These findings collectively show that PAN neural stimulation is mainly contributed by the photoacoustic effect.

Notably under the ns laser with the thermal confinement and stress confinement condition met, many nanoparticles, including Au nanoparticles, can also be photoacoustic ^41^. The photoacoustic properties of these nanostructures have been only applied for photoacoustic imaging. Our work is the first-time demonstration of photoacoustic nanoparticles for neurostimulation. The semiconducting polymer-based PAN provides a new paradigm for neural modulation through offering two important features compared to other photoacoustic agents. First, we compared COMSOL simulation for Au nanoparticles under the ns laser condition at the wavelength of 532 nm wavelength (**Figure S15**) to that for PANs. Under the same laser power, the maximum temperature rise is 40.4 °C on Au nanoparticle surface, compared to 8.4 °C on PAN surface. As it produces less temperature rise avoiding potential thermal toxicity while effectively activating neurons, PAN is of particular interest for neuron stimulation. Second, the semiconducting polymer-based PANs provide an exciting opportunity for non-invasive neural modulation. Our PANs uniquely absorb NIR-II light. Due to its longer wavelength, NIR-II light has been reported to have sufficient penetration depth in highly scattering medium ^23,42,43^. Such wavelength has also been demonstrated to have the capability of penetrating human skull ^20,21^, potentially enabling non-surgical brain stimulation through light excitation. To illustrate the possibility for deep penetration, we embedded PANs in a 5 mm thick brain-mimicking phantom under a skull. We clearly detected ultrasound signals from these PANs by ns laser excitation above the skull using photoacoustic tomography (**Figure S16**). In addition, advances in biophotonics showed that NIR light focusing with approximately 100 µm is possible in brain tissue ^44^. Together with potential development in surgical free targeted delivery of PANs to specific regions of a brain, for example, via ultrasound openings of the blood-brain barrier ^45,46^, PANs promise an opportunity of non-genetic and non-surgical brain modulation in live animals and further in human patients.

## Materials and Method

### Materials

All chemical reagents were purchased from Sigma-Aldrich (MO, USA) unless otherwise stated. The semiconducting polymer was synthesized via palladium-catalyzed C–C cross-coupling techniques and was thoroughly purified to remove any metal residues. The TRPV4 antibody powders were purchased from Aloemone lab and used without further modification.

### Characterization

SEM images were obtained on a Zeiss Supra 55 scanning electron microscope with an accelerating voltage of 3 kV. A sputter coater, Cressington 108, was used to coat a layer of Au/Pd on biological samples for SEM imaging. DLS and zeta potential measurements were performed on a Brookhaven 90plus nano-particle sizer with zeta potential. UV-Vis-NIR spectra were recorded on a Cary 5000 spectrophotometer. FT-IR spectrum was taken on a Bruker Optics Vertex 70v FTIR, equipped with a Hyperion microscope and Silicon Bolometer.

### Synthesis of PAN

Following the previously developed synthesis process ^23^, the polymers were dissolved in THF (1 mg/mL) with surfactant PS-PAA (5 mg/mL) rapidly injected into deionized water (9 mL) under continuous sonication with a microtip-equipped probe sonicator (Branson, W-150) at a power output of 6 W for 30 s. After sonication for an additional 1 min, THF was removed by nitrogen bubbling for 3 hours. The aqueous solution was filtered through a polyethersulfone (PES) syringe driven filter (0.22 µm) and centrifuged three times using a 30 K centrifugal filter unit at 3500 rpm for 15 min. PAN solution was stored in dark at 4 °C for further use.

### Photoacoustic measurement of PAN

The PAN solution (1 mg/mL) was added into a polyurethane capillary tube with two ends fixed with Epoxy. A customized and compact passively Q-switched diode-pumped solid-state laser (1030 nm, 3 ns, 100 μJ, repetition rate of 3.3 KHz, RPMC, Fallon, MO, USA) was used as the excitation source. The laser was connected to an optical fiber through a homemade fiber jumper (SMA-to-SC/PC, ∼81% coupling efficiency). The laser was adjusted to set the output power from the fiber jumper to be approximately 55 mW. The capillary tube was fixed in the water tank. One miniaturized ultrasound transducer with a central frequency of 5 MHz (XMS-310-B, Olympus, Waltham, MA, USA) was used to record the optoacoustic signals. The ultrasonic signal was first amplified by an ultrasonic pre-amplifier (0.2– 40 MHz, 40 dB gain, Model 5678, Olympus, Waltham, MA, USA) and then sent to an oscilloscope (DSO6014A, Agilent Technologies, Santa Clara, CA, USA) to readout. The signal was averaged 20 times. The pressure of the photoacoustic waves generated was calibrated using a hydrophone with a diameter of 2 mm and frequency range of 1-20 MHz (Precision Acoustics Inc., Dorchester, UK). All of the devices were synchronized by the output from the active monitoring photodiode inside the laser.

### Animals

Embryonic day (E) 15-18 rats, obtained from pregnant Sprague−Dawley rats were used for the *in vitro* experiments. Adult (age 14-16 weeks) C57BL/6J mice were used for *in vivo* experiments. All animal care was carried out in accordance with the National Institute of Health Guide for the Care and Use of Laboratory Animals (NIH Publications No. 80−23; revised 1996) and was operated under protocol 17-022 approved by Boston University Animal Care and Use Committee.

### Embryonic neuron culture

The glass-bottom culture dishes used in the embryonic neuron cell cultures were immersed in 0.01% Poly-D-Lysine (Sigma-Aldrich, MO) for overnight at 4 °C and washed in PBS before culture initiation. Primary cortical neurons were obtained from SD rat. Cortices were dissected out from embryonic day 15-18 (E15-18) rats of either sex and digested in 0.05% trypsin/ethylenediaminetetraacetic acid (EDTA, VWR, PA) 15 min at 37 °C and triturated every 5 min. Dissociated cells were washed with and triturated in 10% heat-inactivated fetal bovine serum (FBS, Atlanta Biologicals, GA), 5% heat-inactivated horse serum (HS, Atlanta Biologicals, GA), 2 mM Glutamine-Dulbecco’s Modified Eagle Medium (DMEM, Thermo Fisher Scientific Inc., MA), and cultured in cell culture dishes (100 mm diameter) for 30 min at 37 °C to eliminate glial cells and fibroblasts. The supernatant containing neurons was collected and seeded on poly-D-lysine coated cover glass and incubated in a humidified atmosphere containing 5% CO2 at 37 °C with 10% FBS + 5% HS + 2 mM glutamine DMEM. After 16 h, the medium was replaced with Neurobasal medium (Thermo Fisher Scientific Inc., MA) containing 2% B27 (Thermo Fisher Scientific Inc., MA), 1% N2 (Thermo Fisher Scientific Inc., MA), and 2 mM glutamine (Thermo Fisher Scientific Inc., MA).

### TA microscopy

TA images were obtained as previously described ^25,26,47^. For each TA image, the Z position of the focus was adjusted near the equatorial plane of the neurons so that the soma and neurites were both clearly visualized. The pump (845 nm) and Stokes (1045 nm) powers before the microscope were maintained at 20 and 20 mW, respectively. Both the pump and Stokes beams were linearly polarized. No cell or tissue damage was observed. Images were acquired at 2 μs pixel dwell time.

To quantify the number of PANs that bind onto neurons, the following estimation was used. An effective concentration of bound PANs can be estimated based on the TA intensity in the image and the TA calibration curve. The TA calibration curve was obtained TA intensities obtained from TA images acquired for PAN solutions with known concentrations. The volume occupied by the bound PANs or PAN-TRPV4 was estimated based on the area of bounded PANs on neuronal soma (measured from Image J) and TA focal depth of 1 µm. The molecular weights of PAN and PAN-TRPV4 were estimated based on electrophoresis measurement to be 75 kD and 125 kD. The number of PAN or PAN-TRPV4 was calculated based on the above-mentioned parameters.

### SEM imaging of PAN on neurons

Cells were fixed in glutaraldehyde solution and then rinsed three times in phosphate-buffered saline (PBS, VWR, PA). Subsequently, samples were immersed in OsO_4_ (Sigma-Aldrich, MO) for 40 min and then rinsed 3 times in PBS, followed by dehydration. The dried samples were mounted on aluminum stubs and coated with platinum for imaging.

### Cytotoxicity tests

Neurons were seeded in 96-well plates (1,000 cells/well in 100 µL) and incubated. PAN solution with concentrations of 20, 40, 60 µg/mL was added to the cell culture medium, respectively. Neurons were incubated with PANs or cell culture medium only for 1 hour and 24 hours, respectively. The medium was then removed and washed with PBS. MTT (20 µL, 5 mg/mL) was added to the wells and incubated for 5 h. The cell culture medium was then removed, and dimethylsulfoxide (DMSO, 200 µL) was subsequently added to each well. Finally, the plates were gently shaken for 10 min at room temperature to dissolve all formed precipitates. The absorbance of MTT at 590 nm was measured using a SpectraMax plate reader. Cell viability was expressed as the ratio of the absorbance of the cells incubated with PANs to that of control.

### Synthesis of PAN-TRPV4

PAN solution (1 mL, 20 μg/mL) was combined with 10xPBS (110 μL, pH=7.4), followed with NHS (30 μL, 9 mg/mL) and EDC (50 μL, 9 mg/mL). The mixture was stirred for 1 hour at room temperature in dark. Anti-TRPV4 antibody (5 μL, 1 mg/mL) was added into the mixture and kept stirring at room temperature overnight. The resulted solution was filtered through a polyethersulfone (PES) syringe driven filter (0.22 µm) and centrifuged three times using a 50 K centrifugal filter unit at 3500 rpm for 15 min. PAN-TRPV4 solution was stored in dark at 4 °C for further use.

### Agarose gel electrophoresis

Agarose gel (0.5 %) was prepared using agarose power purchased from Bio-Rad (Certified Molecular Biology Agarose) and immersed in 1× Tris/boric acid/EDTA (TBE) buffer with 0.1% sodium dodecyl sulfate (SDS). The PAN and PAN-TRPV4 solution (0.1 mg/mL, 10 µL) mixed with SDS (1 µL, 10%) was loaded into the wells of the gel. The electrophoresis process was operated on a horizontal electrophoresis system (Mini-Sub Cell GT; electrode spacing 15 cm) for 1 hour at 150 V. The location of nanoparticles was imaged on a white lightbox with a digital camera (Sony α7).

### Immunofluorescence imaging

Cells were fixed with 4% paraformaldehyde for 20 min. After 3 washes, cells were blocked in 5% bovine serum albumin (Sigma-Aldrich, MO) for 30 min and permeabilized with 0.2% Triton X-100 (Sigma-Aldrich, MO). Then cells were incubated with mouse monoclonal anti-TRPV4 (1:1000) antibody for 2 hour, and then with goat anti-mouse secondary antibody (1:1000, 488 nm, Thermo Fisher Scientific Inc., MA) for 1 hour at room temperature. The fluorescence images were taken with a home-built wide-field fluorescence microscope (Olympus).

### In vitro neurostimulation

PAN or PAN-TRPV4 (150 μL, 20 μg/mL) solution was added into the culture medium of GCaMP6f labeled neurons to reach a final concentration of 2 μg/mL. An incubation time of 15 minutes and 1 hour was tested. A Q-switched 1030-nm nanosecond laser (Bright Solution, Inc.) was used. For the CW laser, Cobolt Rumba 1064 nm 500 mW laser was used. The CW laser was gated with a mechanical shutter to control the laser duration. The laser was delivered using an optical fiber (Thor lab) with a diameter of 200 µm and 0.22 NA. During neurostimulation experiments, the fiber was placed 100 µm approximately above the neurons and the illumination area is calculated to be 222.27 µm. Calcium fluorescence imaging was performed on a lab-built wide-field fluorescence microscope. The microscope was based on an Olympus IX71 microscope frame. With a 20× air objective (UPLSAPO20X, 0.75NA, Olympos), illuminated by a 470 nm LED (M470L2, Thorlabs) and a dichroic mirror (DMLP505R, Thorlabs). Image sequences were acquired with a scientific CMOS camera (Zyla 5.5, Andor) at 20 frames per second. The fluorescence intensity analysis and exponential curve fitting were performed using ImageJ (Fiji).

### In vivo injection of PANs

Adult (age 14-16 weeks) C57BL/6J mice were used. Mice were initially anesthetized using 5% isoflurane in oxygen and then placed on a standard stereotaxic frame, maintained with 1.5 to 2 % isoflurane. Toe pinch was used to determine the level of anesthesia throughout the experiments and body temperature was maintained with a heating pad. The hair and skin on the dorsal surface targeted brain regions were trimmed. Craniotomies were made on primary motor cortex based on stereotaxic coordinates using a dental drill and artificial cortical spinal fluid was administrated to immerse the brain. PAN solution with a concentration of 1.0 mg/mL was injected into the primary motor cortex using a quintessential stereotaxic injector (Stoelting Co.) with a speed of 20 nL/min for 30 min. The injected PANs were allowed to diffuse for 1 hour before stimulation experiments. Optical fiber with a diameter of 200 μm was coupled to a micromanipulator to control the illumination position.

### In vivo LFP recording

Electrophysiology was performed using multifunctional fibers with a thick polymer coating as the recording electrodes ^33,34^. The electrodes were positioned with a micromanipulator to the targeted brain region (Siskiyou). Extracellular recordings were acquired using a Multi-Clamp 700B amplifier (Molecular Devices), filtered at 0.1 to 100 Hz, and digitized with an Axon DigiData 1550 digitizer (Molecular Devices).

### EMG recording

EMG was performed using needle electrode inserted subcutaneously and parallel to the forelimb triceps brachii muscle. Reference electrode was inserted in the footpad. A ground electrode was inserted subcutaneously on the trunk and ipsilateral to the stimulation site. EMG signals were acquired using a Multi Clamp 700B amplifier (Molecular Devices), filtered at 1 to 5000 Hz, and digitized with an Axon DigiData 1550 digitizer (Molecular Devices).

### COMSOL simulation

The temperature change on PANs and gold nanoparticles was simulated in COMSOL Multiphysics (COMSOL Inc, USA). The size of the nanoparticles for both PAN and gold nanoparticle is set to be 60 nm. The laser irradiation on a single PAN is assumed to be uniform. The surrounding environment is water. The absorption cross-sections of both PAN and gold nanoparticles are calculated based on Mie scattering theory ^48^. To calculate the temperature change induced by laser irradiation, the heat transfer module was used. The simulation model was validated first by comparing the simulation results to the experimental data reported previously ^12^.

### Photoacoustic tomography

The photoacoustic signals were processed by a high-frequency ultrasound imaging system (Vantage 128, Verasonics Inc). A brain mimicking phantom (10% Gelatin, 1 % formaldehyde and 5% intralipid) was used to mimic the highly scattered brain tissue ^49^. A piece of mouse skull was placed on top of the phantom to test the penetration depth of NIR-II window light. PAN solution with a concentration of 1.5 mg/mL in a transparent polyurethane tube was place under the phantom. A Q-switched Nd:YAG laser (CFR ICE450, Quantel Laser) with 8 ns pulse width and 10 Hz repetition rate was applied as the excitation. The wavelength was set to be 1100 nm and laser power was fixed to be 9 mJ. The laser light was guided to the surface of the mouse skull by a fiber bundle and the photoacoustic signals were detected from the other side of the tissue by a low frequency transducer array (L7-4, PHILIPS/ATL.)

### Data analysis

Calcium images were analyzed using ImageJ. Calcium traces, electrophysiological traces, and EMG recordings were analyzed and plotted using Origin and MATLAB. All statistical analysis was done using Origin. Data shown are mean ± SD. *P* values were calculated based on two sample t-test and defined as: n.s.: non-significant, p > 0.5; *: p < 0.5; **: p < 0.01; ***: p < 0.001.

## Acknowledgement

This work was supported by Boston University Nanotechnology Innovation Center Cross-Disciplinary fellowship for Y.H., R01 NS109794 to J.C. and C.Y., NSF1847436 to X.J.. We thank Margaret O’Connor, Zachary Gardner and Yuan Tian for their assistance in the preparation of primary neuronal cultures.

## Author contributions

C.Y. and J.C. conceived the concept of using PAN for neuromodulation; X. L and J.M. synthesized the semiconducting polymer; Y.H., J. W. and R. C. synthesized PAN; Y. H. and Y. J. designed and performed the experiments; Y. H., Y. J., S. L., H.M., J. C. and C. Y. discussed and analyzed the data; H. M. provided neuron cultures; S. J. and X. J. designed and fabricated the multifunction electrode for in vivo LFP recording; H.Z. performed the COMSOL simulation; Y. H., Y. J. and C. Y. wrote the manuscript. All authors discussed and edited the manuscript. C. Y. and J.C. supervised the project.

