## Supplemental information for "Neural Stimulation in vitro and in vivo by Photoacoustic Nanotransducers"

Corresponding authors:

**List of Figures**

**Figure S1.** FT-IR spectrum confirmed the surface modification of PANs.

**Figure S2.** TA microscopy for imaging PAN solution.

**Figure S3.** SEM images of PANs binds to embryonic neurons soma region.

**Figure S4.** TA images of PANs cultured with neurons and glial cells for 1 hour.

**Figure S5.** Representative curve of the prolonged stimulation with recording time of 60 s.

**Figure S6.** Distribution of decay constants for PAN induced neurostimulation.

**Figure S7.** Spatial distribution of neurostimulation induced by PAN with 3 ms laser duration with synaptic blocker added.

**Figure S8.** Neuromodulation rate as a function of laser duration.

**Figure S9.** Neurostimulation rate as a function of culture time.

**Figure S10.** Digital images of PAN and PAN-TRPV4 solutions and normalized UV-Vis spectrum of PAN and PAN-TRPV4 solution.

**Figure S11.** IF images of TRPV4 channel expression on the neuron membrane labeled with anti-TRPV4 antibody.

**Figure S12.** Distribution of decay constants for PAN-TRPV4 induced neurostimulation.

**Figure S13.** Cross-section image of multifunctional fibers used for neural electrical signal recording.

**Figure S14.** CW laser induced neurostimulation with a laser duration of 2.5 s.

**Figure S15.** Temperature change profile of gold nanoparticles calculated under 3 laser pulses with a pulse width of 3 ns and pulse energy of 1.18 nJ.

**Figure S16.** Photoacoustic/ultrasound image of PAN solution (1.5 mg/mL) in a transparent polyurethane tube placed underneath mouse skull and a brain mimicking phantom with a thickness of 5.5 mm.

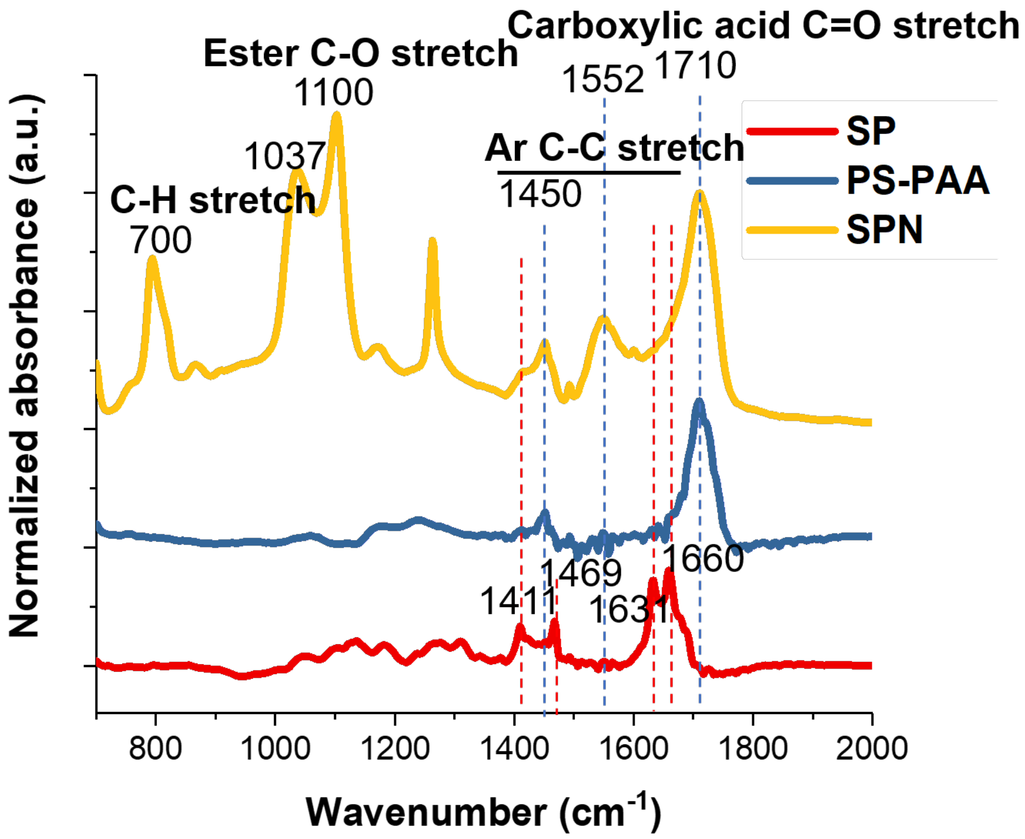


**Figure S1. FT-IR spectrum confirmed the surface modification of PAN.**

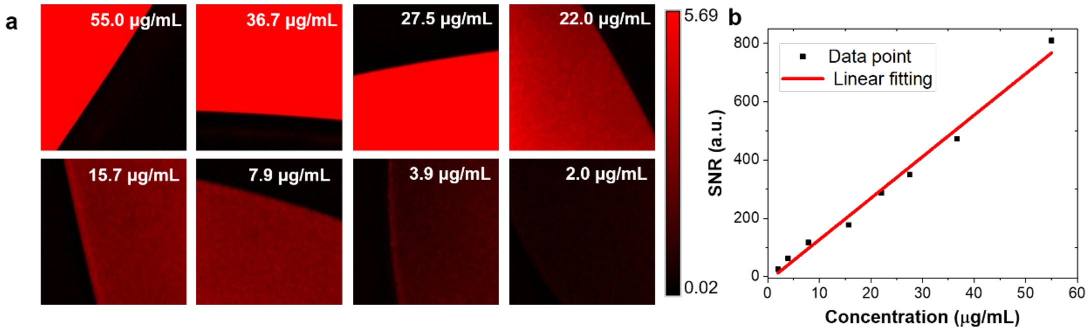


**Figure S2. TA microscopy for imaging SPN solution.** (a)TA imaging of PAN solution with concentration range in 2.0 to 55.0 μg/mL. (b) TA signal-to-noise ratio (SNR) plotted against concentration of PAN solution. Date were fitted linearly.

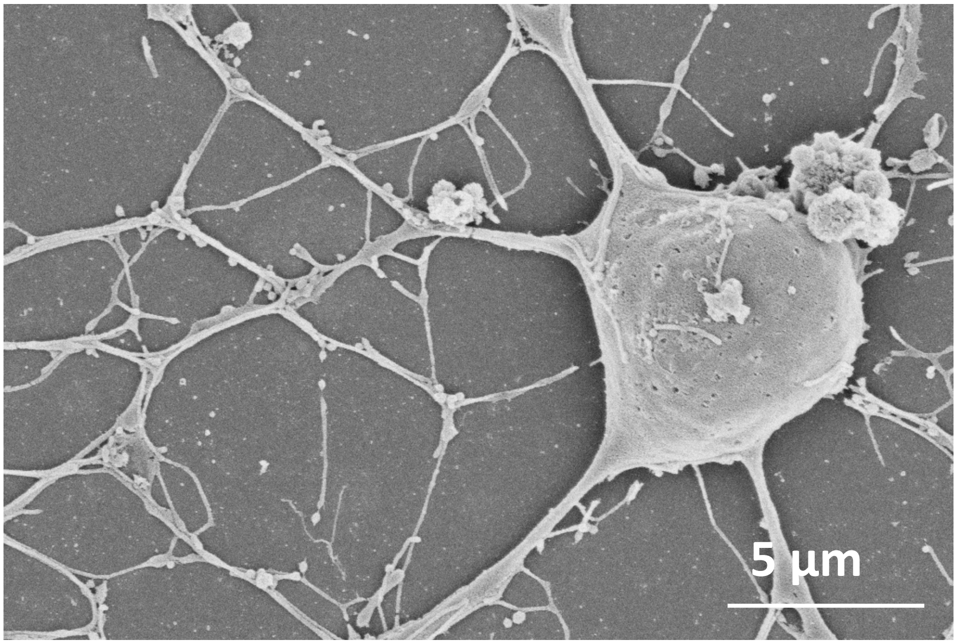


**Figure S3. SEM images of PANs binding to embryonic neurons soma region. Scale bar: 50 μm.**

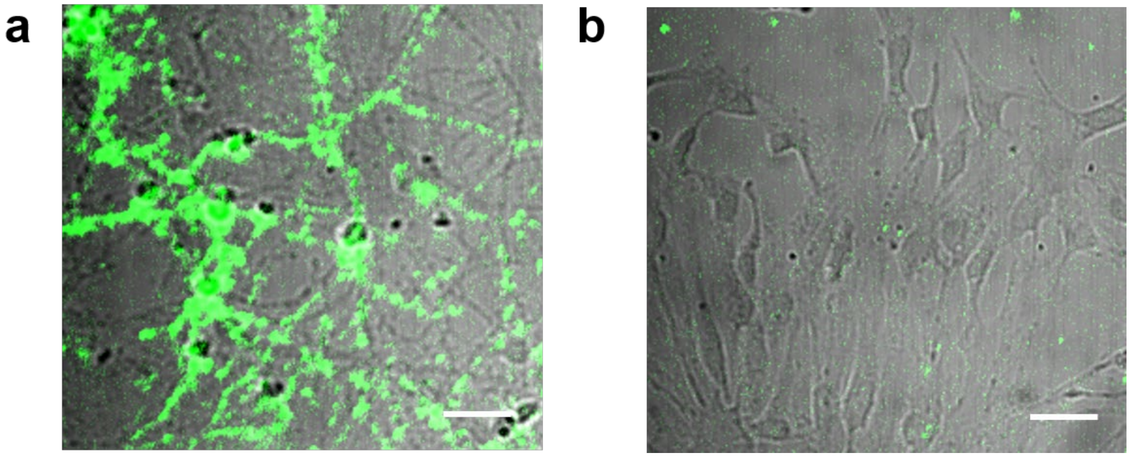


**Figure S4. TA images of PANs cultured with neurons (a) and glial cells (b) for 1 hour. Scale bar: 50 μm.**

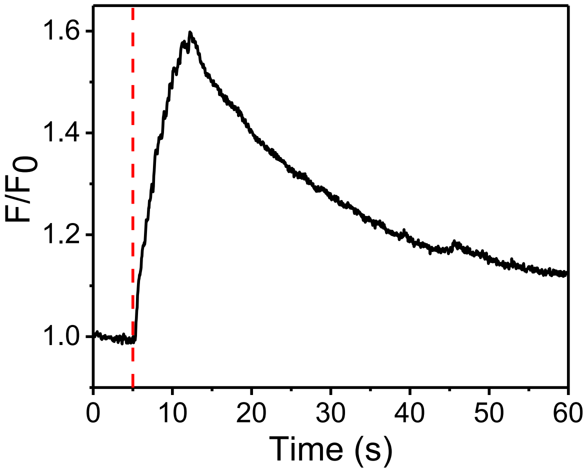


**Figure S5. Representative curve of the prolonged stimulation with recording time of 60 s.**

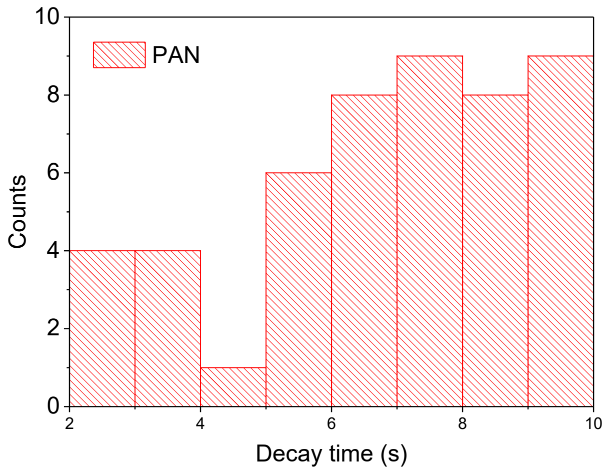


**Figure S6. Distribution of decay constants for PAN induced neurostimulation.**

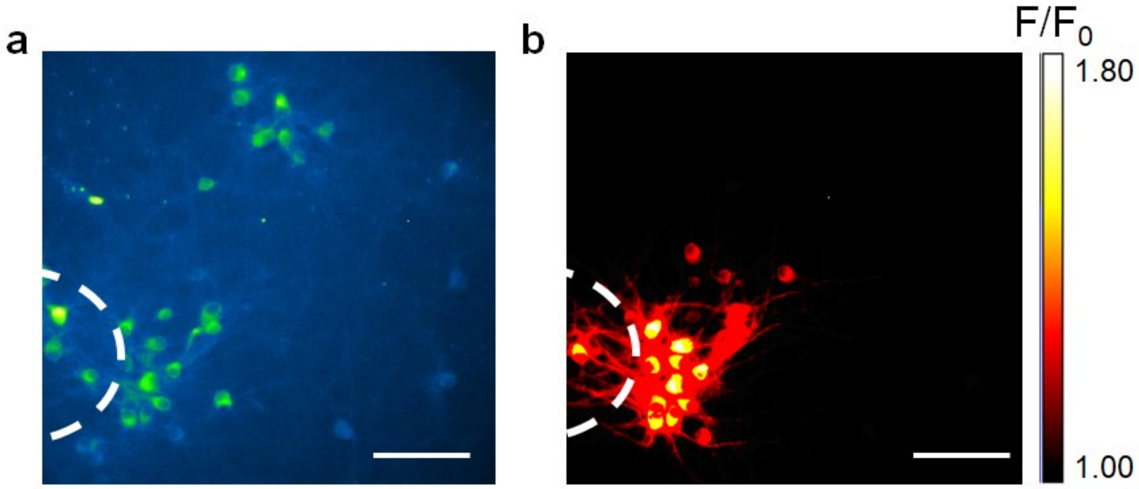


**Figure S7. Spatial distribution of neurostimulation induced by PAN with 3 ms laser duration with synaptic blocker added.** (a) Fluorescence images of neurons before stimulation. (b) $F/F_0$ image 1s after laser onsite. Scale bar: 100 μm. White dash line indicates the position of the optical fiber.

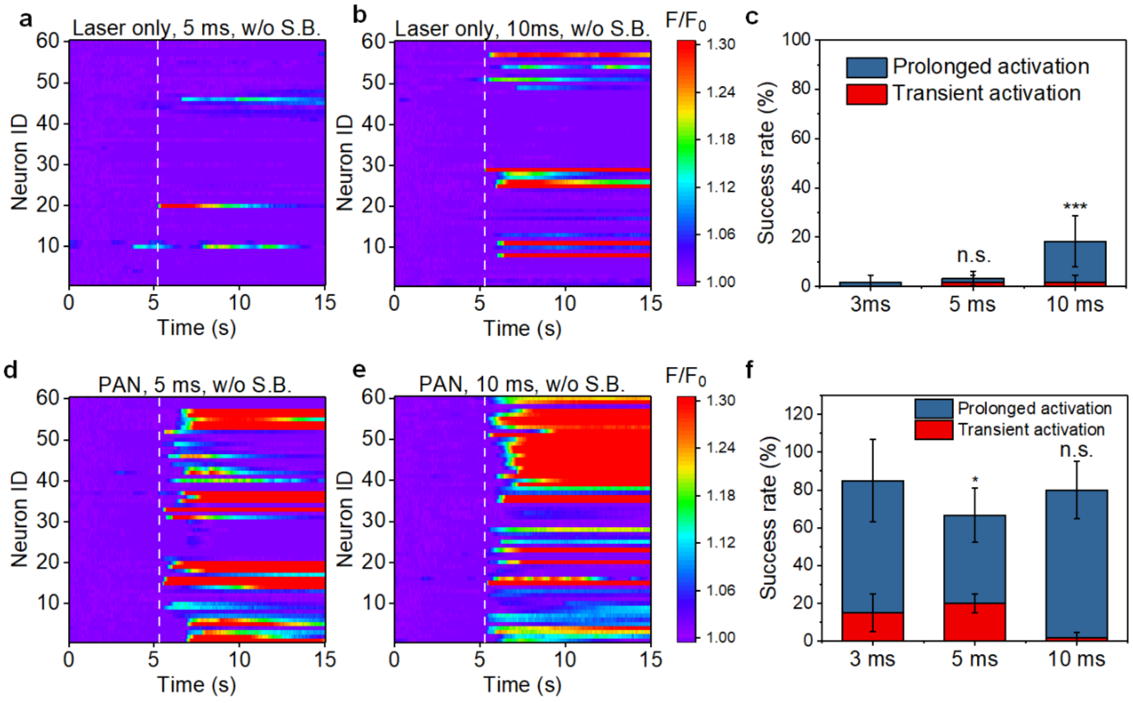


**Figure S8. Neuromodulation rate as a function of laser duration.** (a-b) Colormaps of fluorescence signal change of neurons stimulated by a 1030 nm nanosecond laser only with laser duration of 5 ms (16 pulses, a) and 10 ms (32 pulses, b), respectively. (c) Success rate of nanosecond laser induced neurostimulation. (d-e) Colormaps of fluorescence signal change of neurons cultured with PAN solution for 15 minutes and stimulated by a 1030 nm nanosecond laser only with laser duration of 5 ms (d) and 10 ms (e), respectively. (f) Success rate of PAN induced neurostimulation under different laser durations. All Colormaps were plotted under same dynamic range.

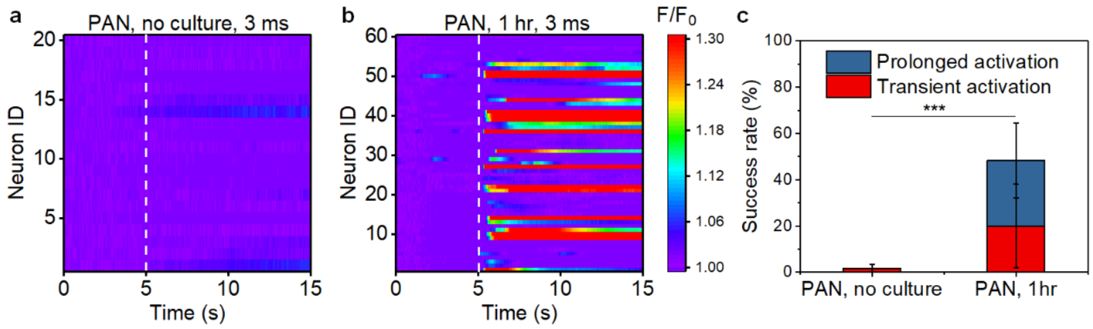


**Figure S9. Neurostimulation rate as a function of culture time.** Colormaps of fluorescence intensity change of no culture (a) and 1-hour culture of PAN (b). (c) Success rate analysis (c).

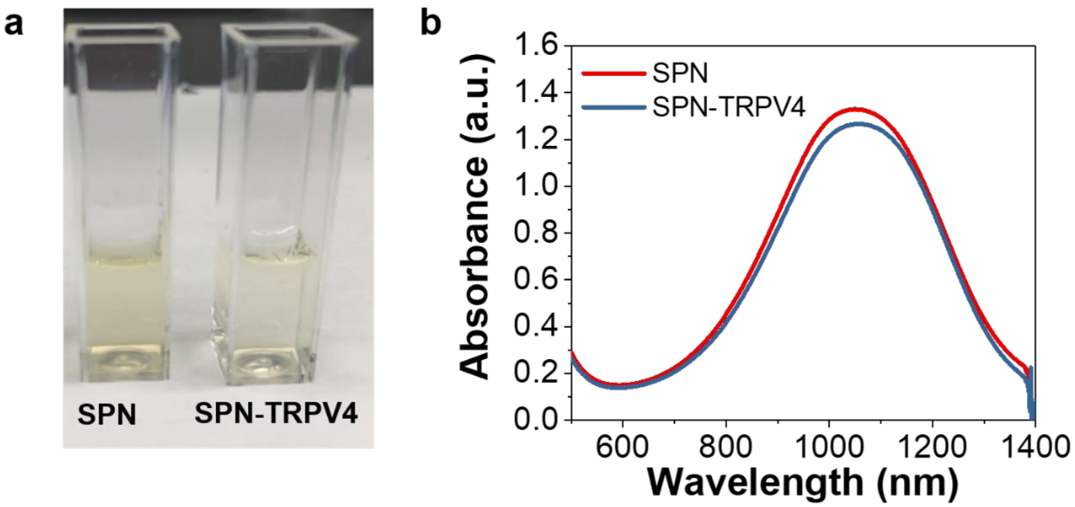


**Figure S10. Digital images of PAN and PAN-TRPV4 solutions with concentration of 20 μg/mL (a). Normalized UV-Vis spectrum of PAN and PAN-TRPV4 solution (b).**

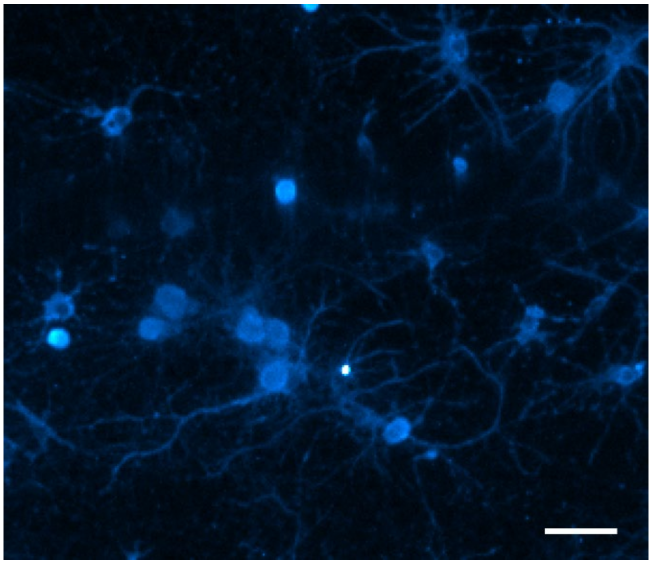

**Figure S11. IF images of TRPV4 channel expression on the neuron membrane labeled with anti-TRPV4 antibody.** Scale bar: 20 μm.

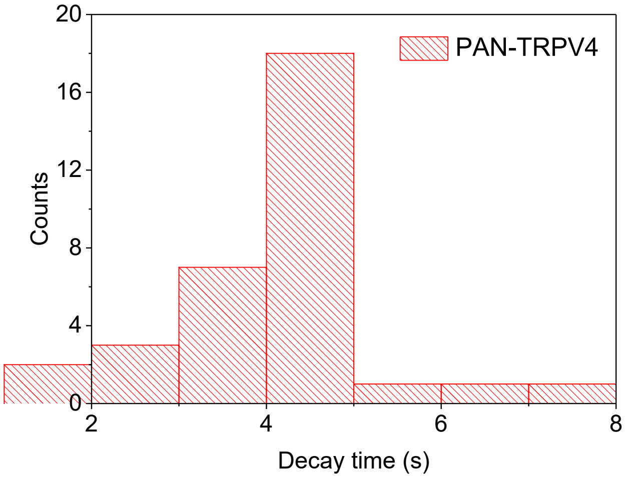


**Figure S12. Distribution of decay constants for PAN-TRPV4 induced neurostimulation.**

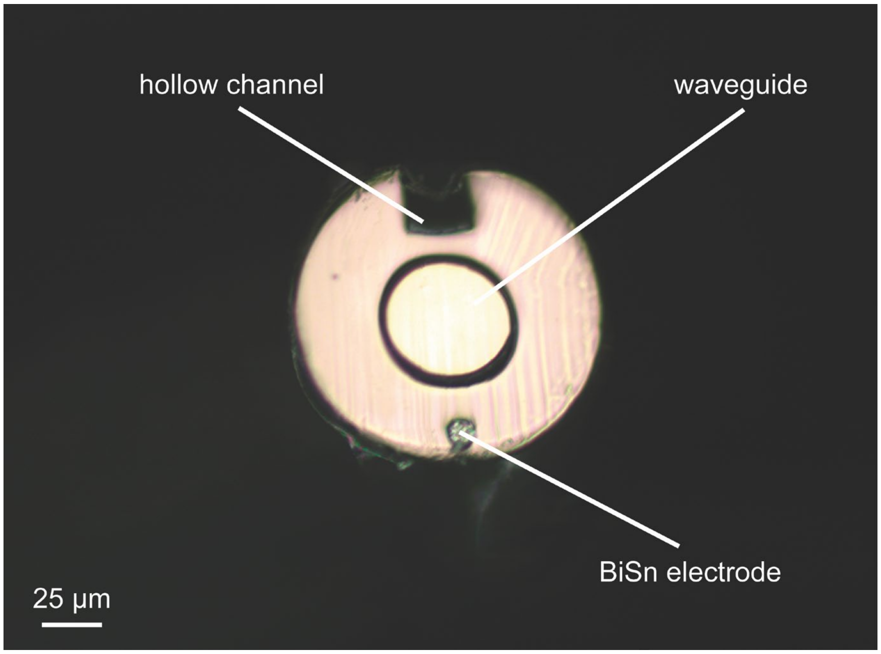


**Figure S13. Cross-section image of multifunctional fibers used for neural electrical signal recording.**

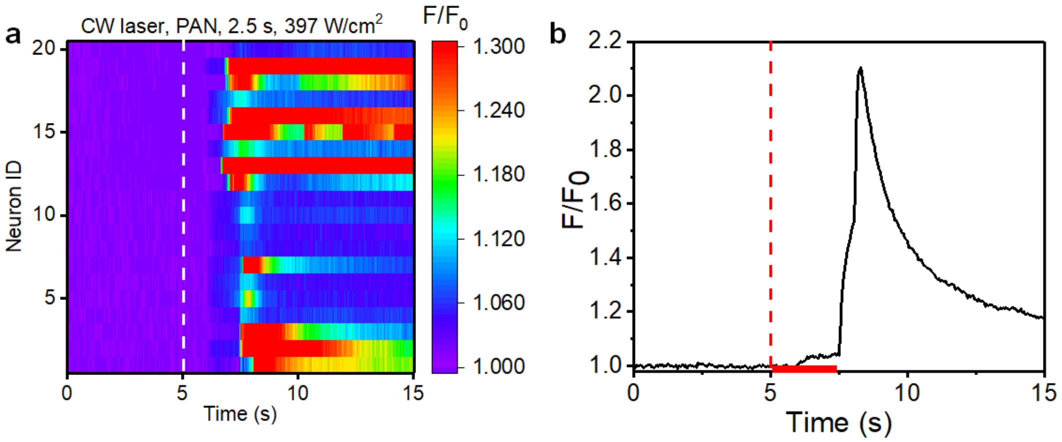


**Figure S14. CW laser induced neurostimulation with a laser duration of 2.5 s.** (a) Colormaps of fluorescence intensity change using CW laser with a 2.5s laser duration. (b) Representative curve of neurostimulation response.

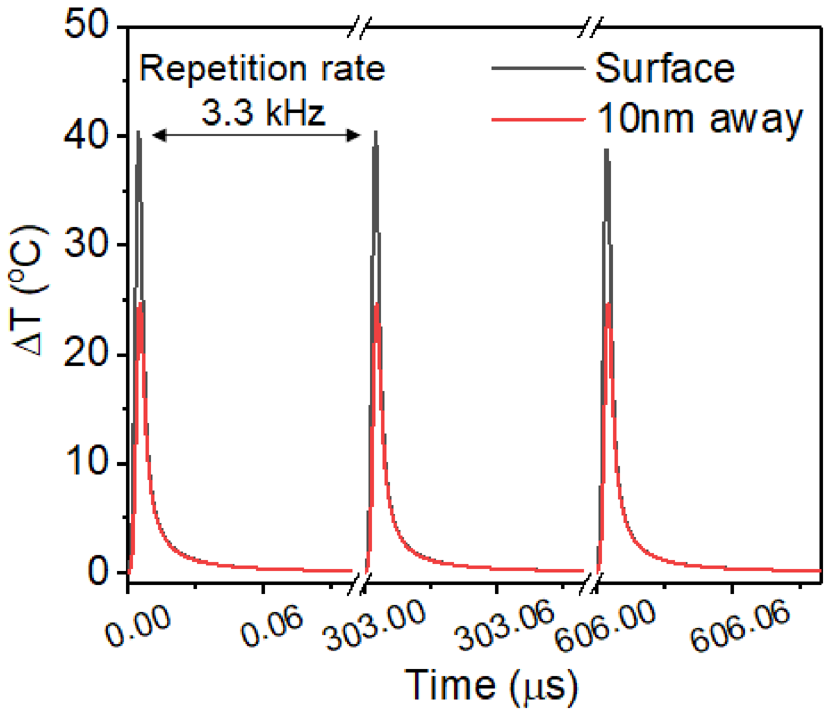


**Figure S15. Temperature change profile of gold nanoparticles calculated under 3 laser pulses with a pulse width of 3ns and pulse energy of 1.18 nJ.**

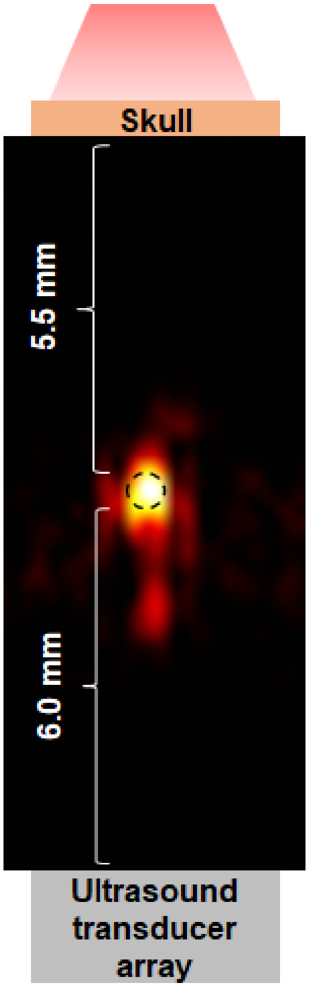


**Figure S16. Photoacoustic/ultrasound image of PAN solution (1.5 mg/mL) in a transparent polyurethane tube (black circle) placed underneath mouse skull and a brain mimicking phantom with a thickness of 5.5 mm.**
